# Intrinsic and extrinsic factors collaborate to activate pharyngeal satellite cells without muscle injury

**DOI:** 10.1101/2020.05.21.108951

**Authors:** Eunhye Kim, Yiming Zhang, Fang Wu, James Allen, Katherine E. Vest, Hyojung J. Choo

**Affiliations:** Department of Cell Biology, School of Medicine, Emory University, Atlanta, GA 30322, USA; Department of Molecular Genetics, Biochemistry and Microbiology, University of Cincinnati College of Medicine, Cincinnati, OH 45267, USA

**Keywords:** Skeletal muscle stem cells, Satellite cells, Pharyngeal muscle, Satellite cell activation, Craniofacial muscle

## Abstract

Satellite cells (SCs), adult muscle stem cells in craniofacial muscles proliferate and differentiate/fuse without injury, unlike quiescent SCs in uninjured limb muscle. However, whether intrinsic or extrinsic factors driving their increased basal activity are largely unknown. We compared SCs from the pharynx, which contains constrictor muscles critical for swallowing, to SCs from limb muscle. Pharyngeal SCs are intrinsically more proliferative and contain higher mitochondrial content relative to limb SCs. Pharyngeal SCs occupy less quiescent microenvironments containing collagen V and pharyngeal muscles provide a distinctive SC niche enriched with neighboring resident macrophages and fibroadipogenic progenitors. Loss of SCs impacts pharyngeal myofiber cross-sectional area and the number of neighboring cells, suggesting that SCs are required to maintain pharyngeal muscle homeostasis and its unique niche. Taken together, this study gives new insights to explain the distinctive SC activity of craniofacial muscles, which may explain their unique susceptibility to various muscular dystrophies.

## 1. Introduction

The pharynx is a muscular passageway of the digestive and respiratory tracts extending from the nasal and oral cavity to the larynx and esophagus. The pharynx contains a group of skeletal muscles that play a critical role in many vital processes such as swallowing, breathing, and speaking. Like other craniofacial muscles, pharyngeal muscles originate from non-segmented cranial mesoderm during vertebrate embryogenesis, while trunk and limb muscles are derived from somites (Mootoosamy & Dietrich, 2002; Noden & Francis-West, 2006). These distinctive embryonic origins are associated with unique transcriptional regulatory networks in myogenic progenitor cells as exemplified by PAX3-dependent limb muscle development and PITX2/TBX1-dependent craniofacial muscle development. However, both early muscle development pathways converge to a common myogenic program that requires expression of myogenic regulatory factors such as MYF5, MYOD and myogenin (Goulding, Lumsden, & Paquette, 1994; Relaix, Rocancourt, Mansouri, & Buckingham, 2005; Shahragim Tajbakhsh, Rocancourt, Cossu, & Buckingham, 1997). While mature craniofacial and limb/trunk muscles are histologically very similar, they are differentially susceptible to muscular dystrophies. For example, extraocular muscles are typically spared in Duchenne muscular dystrophy (Khurana et al., 1995) but are preferentially affected by oculopharyngeal muscular dystrophy (OPMD), a late-onset genetic disorder characterized by progressive dysphagia and ptosis (Victor, Hayes, & Adams, 1962). Thus, the distinct embryonic origins of craniofacial muscles could drive functional consequences in adult muscles.

An important common feature of craniofacial and limb/trunk muscles is presence of muscle specific stem cells termed satellite cells (SCs). SCs are a heterogeneous population of progenitor cells underneath the basal lamina of muscle fibers and are crucial for skeletal muscle regeneration (Lepper, Conway, & Fan, 2009; Mauro, 1961; Sambasivan et al., 2011). Like most other adult stem cells, SCs are quiescent under homeostatic basal physiological conditions. When activated by injury or disease, SCs rapidly re-enter the G_1_ phase of the cell cycle, proliferate as myoblasts, and progress along a defined differentiation program known as myogenesis (Shi & Garry, 2006). The properties of SCs during skeletal myogenesis have been extensively investigated using easily accessible limb muscles, but some groups have expanded studies to SCs in other muscle types including craniofacial muscles. According to these studies, SCs from pharyngeal muscles (Randolph et al., 2015) and extraocular muscles (EOMs (Stuelsatz et al., 2015)) contain a population of activated SCs that chronically proliferate and differentiate into myofibers in the absence of muscle damage. The increased SC activity in craniofacial muscles raises the question of whether their unique biological properties are influenced by cell intrinsic factors or by the specialized microenvironment, known as the niche. Multiple studies have demonstrated that extracellular components like collagen (Baghdadi et al., 2018), diffusible cytokines, and growth factors released from neighboring cells such as resident or infiltrating macrophages and fibroadipogenic progenitors (FAPs) (Evano & Tajbakhsh, 2018) have a major influence on satellite cell activity in limb muscles (Vishwakarma, Rouwkema, Jones, & Karp, 2017). In contrast, very few studies have probed how the unique niche of craniofacial muscles affects SC activity (Formicola, Marazzi, & Sassoon, 2014).

In this study, we compared SCs from pharyngeal and gastrocnemius muscles and discovered distinctive intrinsic attributes of pharyngeal SCs and extrinsic factors of the pharyngeal muscle niche, which may contribute to activate pharyngeal SCs without muscle injury. We demonstrate that pharyngeal SCs are larger, have increased mitochondrial content, and show accelerated *in vitro* proliferation and differentiation compared with gastrocnemius SCs. We also show that pharyngeal SCs secrete factors, which may act in an auto/paracrine manner, to induce increased proliferation of SCs relative to factors secreted from limb SCs. We confirm that pharyngeal muscles are enriched with resident macrophages and FAPs, both known to stimulate proliferation and differentiation of SC in vivo, thus providing a unique niche to the resident SC population. Finally, we used SC ablation experiments to determine the contribution of SC to maintenance of pharyngeal muscle and the pharyngeal muscle niche. These studies provide insight into the unique properties of craniofacial muscles, which may explain the differential susceptibility of these muscles to aging and disease.

## 2. Materials and methods

### Mice

C57BL/6J mice (Jax000664), *Pax7* ^*CreERT2/CreERT2*^ mice (Jax017763), *Rosa* ^*tdTomato/tdTomato*^ (Jax007909), *Rosa-DTA* (Jax009669), *Pax3*^*Cre*^ (Jax005549), *Rosa*^*mT/mG*^ (Jax007576) were purchased from Jackson Laboratories (Bar Harbor, ME; www.jax.org). Three or 12 months old mice were used as noted in figure legend. Homozygous *Pax7* ^*CreERT2/CreERT2*^ male mice were crossed with homozygous *Rosa*^*flox-stop-flox-tdTomato*^ (tdTomato) to obtain *Pax7* ^*CreERT2/+*^; *Rosa* ^*tdTomato/+*^ (*Pax7 Cre*^*ERT2*^*-tdTomato*) mice (Sambasivan et al., 2011). To label satellite cells with red fluorescence (tdTomato), tamoxifen, 1 mg (Sigma-Aldrich, St. Louis, MO) per 10 grams body weight, was injected intraperitoneally once daily for 5 days. Flow cytometry was used to determine the recombination efficiency in *Pax7 Cre*^*ERT2*^*-tdTomato* mice. Quantitative polymerase chain reaction (qPCR) was used to determine the recombination efficiency in *Pax7 Cre*^*ERT2*^*-DTA* mice. Experiments were performed in accordance with approved guidelines and ethical approval from Emory University’s Institutional Animal Care and Use Committee and in compliance with the National Institutes of Health.

### Dissection of Muscle Tissue

Pharyngeal tissue dissection was performed as described previously (Randolph et al., 2015). Briefly, histologic sections included pharyngeal tissue extending from the soft palate caudally to the cranial aspects of the trachea and esophagus. Cross sections were prepared in both transverse and longitudinally for circular outside and longitudinal inside muscles, respectively. For collection and isolation of myogenic cells, the larynx and trachea were excluded from pharyngeal samples. Gastrocnemius muscles were used as control limb muscles.

### Satellite Cell Isolation and Fluorescence Activated Cell Sorting

To obtain purified satellite cells (SCs), primary cells were isolated as described previously with small modifications (Randolph et al., 2015). Briefly, dissected pharyngeal and gastrocnemius muscles were minced and digested using 0.2% collagenase II (Gibco, Carlsbad, California) and 2.5U/ml Dispase II (Gibco, Carlsbad, California) in Dulbecco’s modified Eagle’s medium (DMEM) at 37°C while shaken at 80 rpm for 90 minutes. Digested muscles were then rinsed with same volume of Ham’s F10 media containing 20% FBS and 100 µg/ml penicillin/streptomycin (P/S). Then, mononucleated cells were collected using 70 µm cell strainer (Thermo Fisher Scientific, Waltham, MA). To facilitate rapid isolation of a pure pharyngeal and hind limb SCs, we used an antibody-free fluorescence based lineage labeling strategy, whereby Pax7 positive SCs are marked with a red fluorescence, tdTomato, upon tamoxifen-mediated Cre recombinase activation. Fluorescence-activated cell sorting (FACS) was performed using a BD FACSAria II cell sorter (Becton-Dickinson, http://www.bd.com, Franklin Lakes, NJ) at the Emory University School of Medicine Core Facility for Flow Cytometry. Analyses of flow cytometry data were performed using FACSDiva (BD version 8.0.1) and FCS Express 6 Flow. FACS-purified SCs were plated at 500 cells per well in 48-well plate coated with Matrigel (Corning Life Sciences, New York, NY, Ca No. 354277) and cultured for five days in Ham’s F10 media (Hyclone, Pittsburgh, PA; www.gelifesciences.com) containing 20% FBS and 25 ng/mL FGF2 (PeproTech, Rocky Hill, NJ). In Supplementary Figure 1, we used antibody strategy to isolate pharyngeal and hindlimb SCs. Cells were labeled using the following antibodies: 1:400 CD31-PE (clone 390; eBiosciences, San Diego, CA http://www.ebioscience.com/), 1:400 CD45-PE (clone 30-F11; BD Biosciences, www.bdbiosciences.com, San Jose, CA), 1:4000 Sca-1-PE-Cy7 (BD Biosciences, clone D7; www.ablab.ca, Vancouver, Canada), 1:500 a7-integrin-APC (Clone R2F2; AbLab). Dead cells were excluded by 5 µg/ml propidium iodide (PI) staining. SCs were collected according to the following sorting criteria: PI^-^/ CD31^-^/CD45^-^/Sca1^-^/Intergrin7α^+^.

### In Vivo Cell Proliferation Assays by Flow Cytometry

To compare the proliferative abilities of SCs in pharyngeal and hindlimb muscles *in vivo, Bromo-2’-deoxyuridine (BrdU)* assays were performed. Three-month-old C57BL/6 male mice were injected with 10 µg BrdU (Sigma-Aldrich, St. Louis, MO; www.sigmaaldrich.com)/gram body weight intraperitoneally every 12 hours for 2 days before sacrifice. Muscles were dissected and digested as described above. To assess proliferation, isolated mononucleated cells from pharyngeal or gastrocnemius muscles were immunostained with the following antibodies: 1:400 CD31-PE (clone 390; eBiosciences, San Diego, CA http://www.ebioscience.com/), 1:400 CD45-PE (clone 30-F11; BD Biosciences, www.bdbiosciences.com, San Jose, CA), 1:4000 Sca-1-PE-Cy7 (BD Biosciences, clone D7; www.ablab.ca, Vancouver, Canada), 1:500 a7-integrin-APC (Clone R2F2; AbLab). Subsequently cells were labeled for BrdU using a FITC-BrdU flow kit in accordance with the manufacturer’s instructions (BD Biosciences, www.bdbiosciences.com, San Jose, CA). Proliferating SCs were collected according to the following sorting criteria: CD31^-^/CD45^-^/Sca1^-^/Intergrin7α^+^/BrdU^+^.

### Fusion index and nuclei number analysis

For fusion assay, SCs were cultured for 10 days to induce spontaneous differentiation (Stuelsatz et al., 2015). Cells were fixed in 2% formaldehyde in PBS for 10 min at room temperature and stained with Phalloidin-iFluor 594 (abcam, ab176757) for 30 minutes at room temperature. Nuclei were then stained with 4’,6-diamidino-2-phenylindole (DAPI) and cells were mounted with Vectashield (Vector Labs, www.vectorlabs.com, Burlingame, CA). Myoblast fusion was quantified by counting myonuclei in myotubes. Fusion index was calculated as the percentage of nuclei of myotubes with two or more nuclei relative to the total number of nuclei in the images. We randomly collected 10 images for each line.

### MitoTracker staining

Pharyngeal and gastrocnemius muscles were dissected from *Pax7 Cre*^*ERT*^*-tdTomato* mice, digested into mononuclear cells and sorted using flow cytometry. Isolated cells were incubated with 50 nM MitoTracker® Green FM (Life Technologies, catalog number: M-7514) at 37°C for 30 min. The cells were washed twice prior to analysis using the FACS LSR II flow cytometer or by fluorescence microscopy.

### Transwell cultures of satellite cells (SCs) and myogenic progenitor cells (MPCs)

To provide continuous cytokine delivery, we used a co-culture system of sorted tdTomato^+^ (PAX7^+^) SCs from pharyngeal or gastrocnemius muscles as donor cells and expended gastrocnemius myogenic progenitor cells (MPCs) as recipient cells. PAX7^+^ satellite cells from pharyngeal or gastrocnemius muscles settled in the micro-wells at 500 cells/well in the Matrigel-coated 24 well plate. Gastrocnemius MPCs were labelled with 5 μM CellTracker™ Green CMFDA (Life Technologies, C7025) at 37°C for 45 minutes, seeded at 1 × 10^4^ cells/mL in a collagen-coated permeable transwell insert (Corning #3413, Transwell® with 0.4 μm Pore Polyester Membrane Insert), and cultured using MPC culture medium for 1 day to completely adhere. Two days after sorting, transwell inserts (containing MPC) were placed on the micro-wells containing sorted SCs for co-culture and were placed on blank micro-wells (without SCs) as a control. Culture medium was exchanged every other day. On day 4 of co-culture, the number of green fluorescence positive MPCs in the insert was counted and the ratio of cell growth was normalized to MPC number in the blank wells.

### Gene expression analysis by real-time qPCR

The gastrocnemius and pharyngeal MPCs and muscles were analyzed for the expression of related markers by comparative real-time qPCR. Total RNA from samples was extracted using QIAamp RNA blood mini kit (Qiagen, Hilden, Germany) according to the manufacturer’s instructions. Isolated RNA (250 ng) was reverse transcribed into complementary DNA (cDNA) using qScript™ cDNA SuperMix (Quanta Biosciences, Gaithersburg, MD) and then analyzed by real-time qPCR. Amplification of cDNA was performed using Power SYBR® Green Master Mix (Applied Biosystems, Waltham, MA) and 2.5 μM of each primer. All primer sequences are listed in Supplementary Table 1. PCR reactions were performed for 35 cycles under the following conditions: denaturation at 95°C for 15 sec and annealing + extension at 60°C for 1 min. Quantitative levels for all genes were normalized to endogenous *Hprt* expression. Fold change of gene expression was determined using the ΔΔCt method (Livak & Schmittgen, 2001).

### Immunohistochemistry/Immunofluorescence

Immunohistochemistry/Immunofluorescence was performed as follows: sections were incubated with blocking buffer (5% goat serum, 5% donkey serum, 0.5% BSA, 0.25% Triton-X 100 in PBS) for 1 hour and then labeled with primary antibodies (Supplementary Table 2) or isotype controls overnight at 4°C in blocking buffer. The following day, sections were washed three times with washing buffer (0.2% Tween-20 in PBS) and incubated with fluorescence probe-conjugated secondary antibodies for 1 hour at room temperature. We used mannose receptor-1 (CD206) as a marker for resident M2 macrophages and platelet-derived growth factor receptor α (PDGFRα) as a marker for FAPs. The TSA Green kit (Tyramide Signal Amplification; Perkin Elmer, www.perkinelmer.com, Waltham, MA) was used for CD206 and PFGFRα staining to enhance the immunostaining signal, after 1 hour incubation with biotinylated goat-anti-mouse F(ab’)2 IgG fragments (2.5 µg/ml). Nuclei were then stained with DAPI and mounted using Vectashield (Vector Labs, www.vectorlabs.com, Burlingame, CA).

### Statistical Analyses

Statistical analysis was performed using Prism 8.0. Results are expressed as the mean ± standard error of the mean (SEM). Experiments were repeated at least three times unless a different number of repeats is stated in the legend. Statistical testing was performed using the unpaired two-tailed Student’s t-test or ANOVA as stated in the figure legends. p < 0.05 was considered statistically significant. Statistical method, p-values, and sample numbers are indicated in the figure legends.

## 3. Results

### 3.1. Anatomical structure and embryonic origins of pharyngeal muscles

Based on location, the pharynx is separated into three major sections: the nasopharynx (NP), oropharynx (OP), and laryngopharynx (LP). The muscles of pharynx consist of a circular outer layer and a longitudinal inner layer (Fig 1A) (Randolph et al., 2014). The inner layer of the pharyngeal wall is comprised of three paired muscles known as the palatopharyngeus (PP, (Fig 1B-a)), the stylopharyngeus, and the salpingopharyngeus. The outer layer of the pharynx is comprised of three pharyngeal constrictor (PC) muscles: superior (SPC), middle (MPC), and inferior (IPC) (Fig 1A). The IPC is particularly important for swallowing and consists of two muscles, the thyropharyngeus (TP) and cricopharyngeus (CP), which form a sphincter at the transition from the pharynx to the esophagus (Fig 1B-b). During swallowing, the successive contraction of pharyngeal constrictor muscles is required to constrict the pharyngeal lumen to propel the bolus downward to the CP, which is vital to the efficient transfer of the bolus to the esophagus (Cook, 1993). Here, we focused on IPC muscles as CP muscles are involved in several types of pharyngeal pathologies including cricopharyngeal spasm (Búa, Olsson, Westin, Rydell, & Ekberg, 2015) and oculopharyngeal muscular dystrophy (Gómez-Torres et al., 2012).

**Figure 1.**
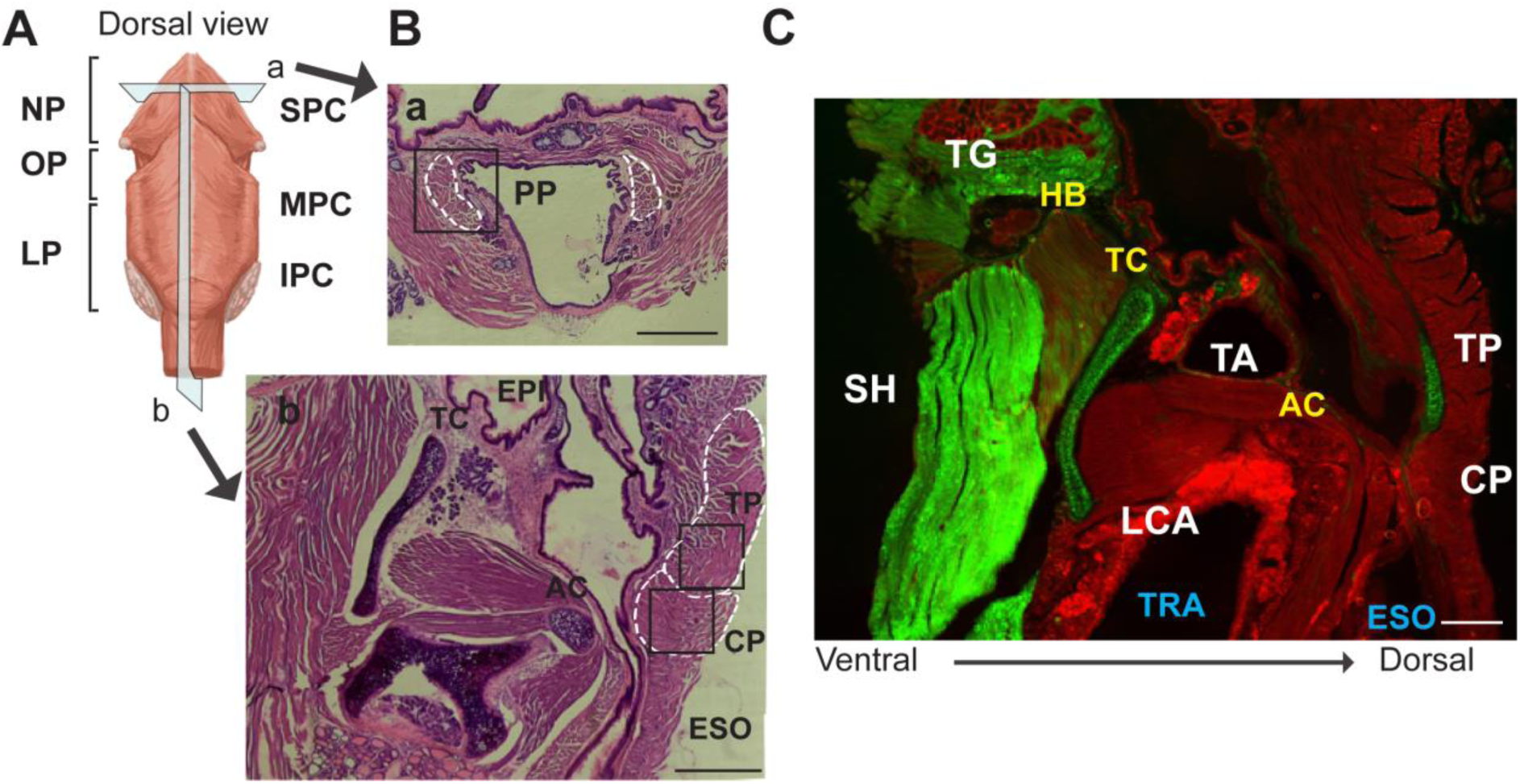
Anatomy and embryonic origins of pharyngeal muscles. A. Illustration of the outer skeletal muscles surrounding the nasopharynx (NP), oropharynx (OP) and laryngopharynx (LP). B. Representative histological images of transverse (a, upper) and longitudinal sections (b, bottom) in pharyngeal muscles. Scale bars = 330 µm. C. Representative longitudinal section of larynx and pharynx expressing PAX3 lineage-derived muscles (green) and non-PAX3 lineage-derived muscles (red) from 20 week old *Pax3*^*Cre/+*^*/mTmG* mice. Abbreviations: superior pharyngeal constrictor (SPC); middle pharyngeal constrictor (MPC); inferior pharyngeal constrictor (IPC); thyropharyngeus (TP); cricopharyngeus (CP); palatopharyngeus (PP); epiglottis (EPI); thyroid cartilage (THY); esophagus (ESO); trachea (TRA); tongue (TG); sternohyoid (SH); hyoid bone (HB); thyroid cartilage (TC); arytenoid cartilage (AC); lateral cricoarythenoid (LCA).

The PAX3 transcription factor is a critical upstream regulator of somitic myogenesis during skeletal muscle development, but it is not expressed in developing craniofacial muscles including pharyngeal muscles (McLoon, Thorstenson, Solomon, & Lewis, 2007; Shahragim Tajbakhsh et al., 1997). To confirm that the PAX3 lineage does not contribute to pharyngeal muscle development, we performed PAX3 lineage tracing using *Pax3*^*Cre/+*^*-mTmG* mice, which label all PAX3 lineage-derived cells with membrane targeted green fluorescent protein (GFP, mG) and non-PAX3 lineage-derived cells with membrane targeted red fluorescent protein, tdTomato (mT) (W. Liu et al., 2013). Similar to extraocular muscles of *Pax3*^*Cre/+*^*-mTmG* mice (Stuelsatz et al., 2015), the TP and CP muscles showed red fluorescence without GFP expression, confirming that they do not originate from PAX3 expressing embryonal progenitors (Fig 1C). Intrinsic laryngeal muscles including thyroarytenoid (TA) and lateral cricoarythenoid (LCA) also expressed tdTomato, indicating that they are non-PAX3 derived muscles. On the other hand, extrinsic laryngeal muscles, such as sternohyoid (SH), and tongue muscles (TG) showed green fluorescence expression, suggesting that the PAX3 lineages contribute to both muscles (Dong et al., 2006; Harel et al., 2009).

Adult SCs are distinguished by expression of the paired-box/homeodomain transcription factor PAX7, which is expressed during quiescence and early activation of SCs and plays a key role in maintenance of self-renewed SCs (Bosnakovski et al., 2008). To investigate the SCs in craniofacial muscles, we used a genetically engineered, tamoxifen-inducible *Pax7 Cre*^*ERT2*^*-tdTomato* mouse, which labels all PAX7 lineage-derived cells with red fluorescent protein (tdTomato). After tamoxifen injection, we observed tdTomato-labeled SCs in sectioned TP, CP, and gastrocnemius (GA, limb) muscles (Fig 2A). The number of SCs in CP muscles was significantly higher than the number of SCs in GA and TP muscles (Fig 2B). Unexpectedly, we also detected significantly increased number of cellular protrusions (filopodia) in SCs from CP muscles relative to SCs in GA muscles (Fig 2C and D). This result is consistent with a previously published study that showed extensive filopodia in extraocular muscles (Verma, Fitzpatrick, & McLoon, 2017), but the role of filopodia in SC function is unknown.

**Figure 2.**
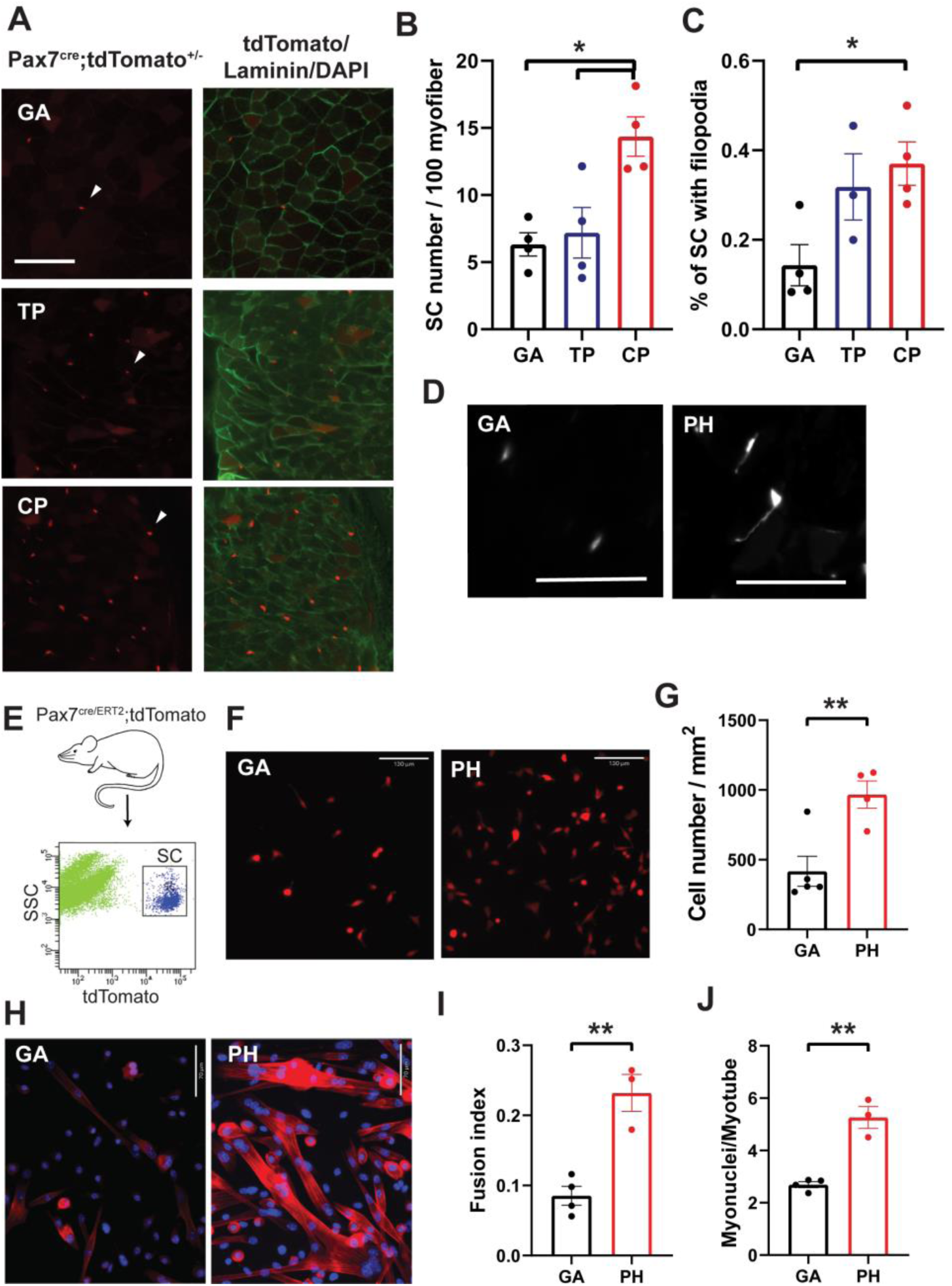
Sorted pharyngeal satellite cells show a high level of proliferation and differentiation. A. Representative cross-section expressing PAX7^+^ SCs (red) in gastrocnemius (GA), thyropharyngeus (TP) and cricopharyngeus (CP) muscles from 3 months old *Pax7 Cre*^*ERT2*^*-tdTomato* mouse. White arrow heads indicate examples of PAX7^+^ tdTomato expressing SCs. Basal lamina was immunostained with anti-Laminin antibody (green). B, C. Quantified numbers of SCs per 100 myofibers (B) and quantified percentages of SCs with filopodia (C) in gastrocnemius and pharyngeal muscles from 3 month old *Pax7 Cre*^*ERT2*^*-tdTomato* mouse. N = 3 or 4. Data were analyzed by 1-way ANOVA. D. Representative image of filopodia in gastrocnemius (GA) and pharyngeal (PH) SCs. Scale bars = 50 µm.E. Scheme of flow cytometry gating strategy for SC isolation using *Pax7 Cre*^*ERT2*^*-tdTomato* mouse. F. Representative image of 5-day cultured myogenic progenitor cells derived from gastrocnemius and pharyngeal SCs of *Pax7 Cre*^*ERT2*^*-tdTomato* mouse. Scale bars = 130µm. G. Analysis of cell number/mm^2^ in gastrocnemius and pharyngeal myogenic progenitor cells derived from SCs of 3 month old mice. n = 4. Statistical significance was determined by Student’s t-test. H. Representative image of gastrocnemius (GA) and pharyngeal (PH) myogenic progenitor cells derived from sorted SCs of *Pax7 Cre*^*ERT2*^*-tdTomato* mice after 10 days of culture. Scale bars = 70 µm. I. Quantified fusion index at 10 days after culture. Fusion index was calculated as the percentage of total nuclei that resided in cells containing 2 or more nuclei. n = 3 or 4. Statistical significance was determined by Student’s t-test. J. Number of myonuclei per myotube at 10 days after culture. n = 3 or 4. Statistical significance was determined by Student’s t-test. For all graphs, the value represents mean ± SEM. Asterisks indicate statistical significance (*p<0.05 and **p<0.01).

### 3.2. Pharyngeal satellite cells are more proliferative and differentiative than gastrocnemius satellite cells in vitro

Pharyngeal SCs are highly proliferative in vivo relative to hindlimb SCs as demonstrated by bromodeoxyuridine labeling and flow cytometric analysis using a known satellite cell gating strategy (CD31^-^/CD45^-^/Sca1^-^/Integrin α7^+^/BrdU^+^) (Fig EV1A) (Randolph et al., 2015). To exclude in vivo niche effects on satellite cell proliferation, we sorted by tdTomato signal equal numbers of pharyngeal and gastrocnemius SCs from *Pax7 Cre*^*ERT2*^*-tdTomato* mice and cultured them for 5 days (Fig 2E and 2F). After 5 days, we detected twice the number of cells in wells containing pharyngeal SCs than those containing SCs from gastrocnemius (Fig 2G). To investigate the differentiation potential of pharyngeal SC, we cultured sorted satellite cells for 10 days to induce spontaneous differentiation (Stuelsatz et al., 2015). The cultured pharyngeal SCs consistently exhibited an increased differentiation at day 10 (Fig 2H) along with an increased fusion index and increased number of myonuclei per myotube compared to the gastrocnemius SCs (Fig 2I and 2J). These results indicate that pharyngeal SCs retain highly proliferative and differentiative properties compared to the limb muscles in the absence of in vivo niche factors, suggesting that cell-intrinsic factors contribute to the unique properties of pharyngeal SCs.

### 3.3. Pharyngeal satellite cells are larger and contain elevated mitochondrial content

When analyzing gastrocnemius and pharyngeal SCs by flow cytometry, two light-scattering parameters, the forward scatter (FSC; cell size) and side scatter (SSC; intracellular granularity and complexity), were also measured. We noticed that FSC values for pharyngeal SCs were 10% higher as compared to gastrocnemius SCs (Fig 3A), suggesting that pharyngeal SCs are larger. The increased cell size and rapid proliferation of pharyngeal SCs resemble properties of the G_Alert_ state that exists in SCs of the contralateral muscles after induced muscle injury mice (Rodgers et al., 2014). Given that G_Alert_ SCs also have increased mitochondrial content and activity, we hypothesized that pharyngeal SCs also have more mitochondria. We stained mitochondria in freshly-isolated SCs and detected increased green fluorescence by microscopy (Fig 3B) and increased relative fluorescence units (RFU) of MitoTracker Green (MTG) by flow cytometry (Fig 3C and 3D) in pharyngeal SCs relative to SCs from gastrocnemius muscle. We observed that the pharyngeal SCs with higher MTG signal (P1 gate of Fig 3D) were larger (increased FCS values) than the pharyngeal SCs with lower MTG signal (Fig 3E). Approximately 12% of pharyngeal SCs had increased cell size and higher mitochondria contents compared to less than 1% of gastrocnemius SCs (Fig 3F). This result indicates that pharyngeal SCs contain increased mitochondrial mass compared with gastrocnemius SCs, which may provide energy and metabolites for early activation and higher proliferation of pharyngeal SCs.

**Figure 3.**
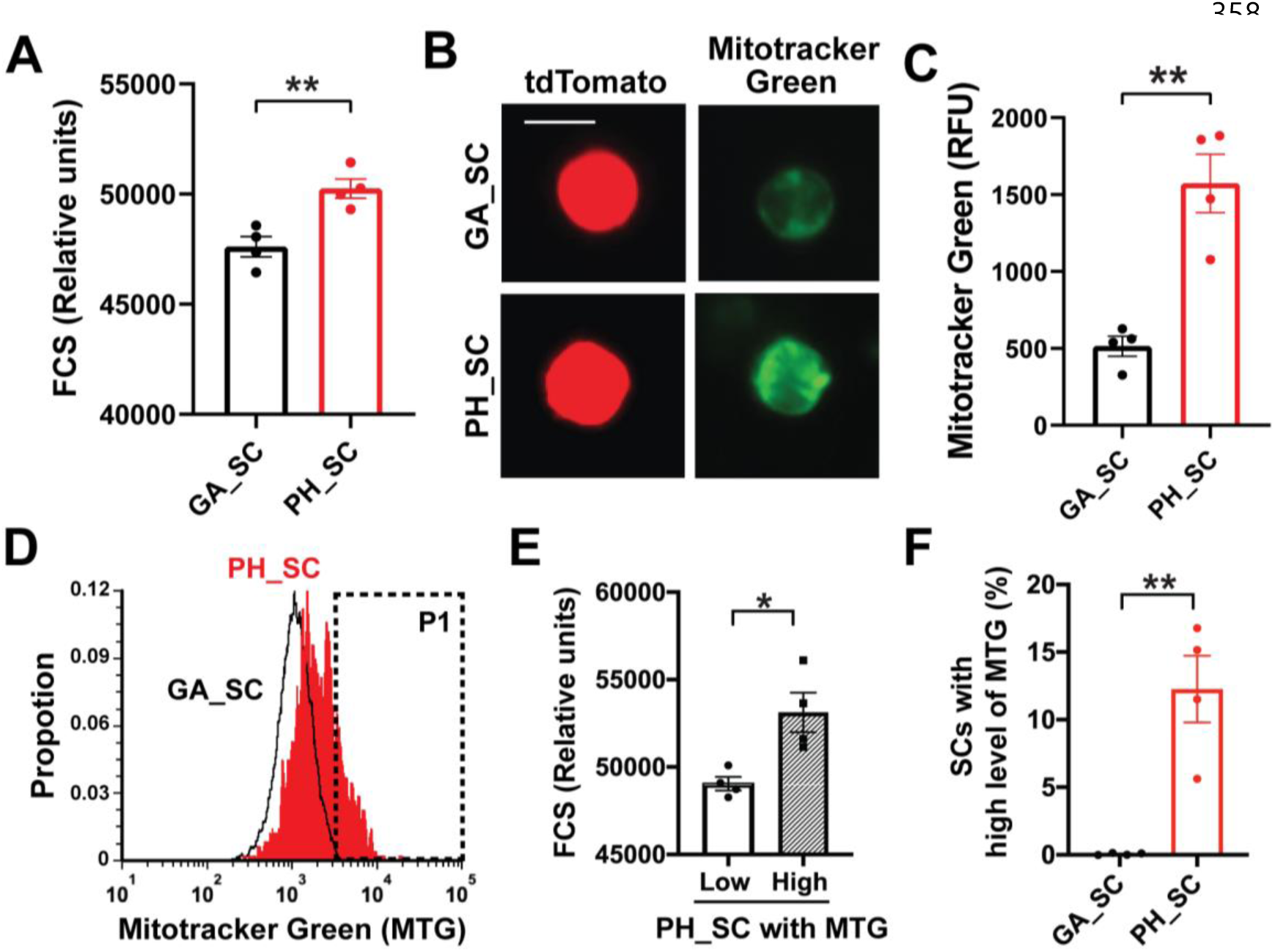
Pharyngeal satellite cells contain more mitochondrial contents. A. Quantified forward scatter (FSC) values for pharyngeal (PH_SC) and gastrocnemius SCs (GA_SC) as determined by flow cytometry. n=4. B. Microscopic fluorescence images showing tdToamto expressing SCs from gastrocnemius or pharyngeal muscles mitochondria stained with MitoTracker Green. Scale bars = 10 µm. C. Representative flow cytometry histogram of MitoTracker Green fluorescence levels of both gastrocnemius (black line) and pharyngeal (red line) SCs. The P1 gate indicates the SC population with high levels of MitoTracker Green. D. Quantitated MitoTracker Green (MTG) relative fluorescence units (y-axis, RFU) of gastrocnemius (black line) or pharyngeal (red line) SCs. n=4. E. Quantified forward scatter (FSC) values for pharyngeal SCs (PH_SC) with low and high MitoTracker Green (MTG) as determined by flow cytometry. n=4. F. Percentage of SCs with high levels of MitoTracker Green (MTG) in pharyngeal (PH_SC) and gastrocnemius SCs (GA_SC). n=4. For all graphs, the value represents mean ± SEM. Statistical significance was determined by Student’s t-test. Asterisks indicate statistical significance (**P<0.01).

### 3.4. Pharyngeal satellite cells secrete proliferation enhancing factors

The increased proliferation of pharyngeal SCs in vitro (Fig 2G) suggests that pharyngeal SCs may secrete pro-proliferative factors. Previous microarray analysis revealed that pharyngeal SCs contain higher levels of RNAs encoding secreted factors that induce cellular proliferation and immune cell infiltration as compared limb SCs (Randolph et al., 2015). To investigate the effect of secreted auto/paracrine factors from gastrocnemius and pharyngeal SCs on proliferation, we sorted PAX7^+^ SCs from *Pax7 Cre*^*ERT2*^*-tdTomato* mice and seeded SCs onto the bottom wells of transwell system. Identical batches of gastrocnemius SCs were seeded onto the upper transwell inserts above either gastrocnemius or pharyngeal SCs and cultured for 48 hours (Fig 4A). We quantified proliferation in the top transwells by counting the number of gastrocnemius SCs dyed with Cell Tracker Green CMFDA (Fig 4B). Compared to controls (blank bottom wells), the number of cells was dramatically increased in both gastrocnemius and pharyngeal SC co-cultured groups (Fig 4C). Importantly, proliferation in the pharyngeal SC group was significantly increased relative to the gastrocnemius SC group. This result suggests that pharyngeal SCs secrete more pro-proliferative factors or fewer anti-proliferative factors during *in vitro* culture compared to gastrocnemius SCs.

**Figure 4.**
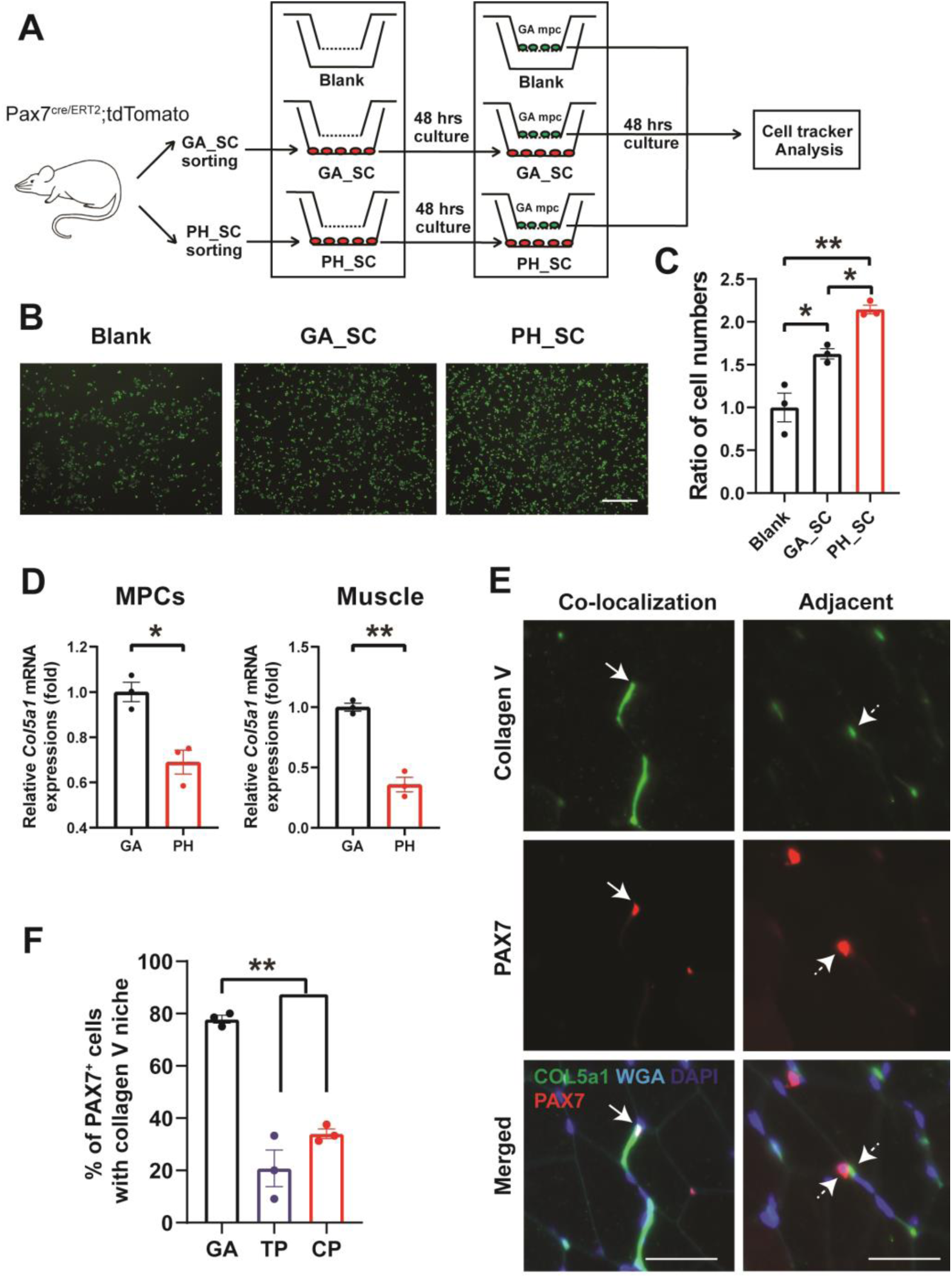
Pharyngeal satellite cells secrete pro-proliferating factors and partially connect with quiescent niche. A. Schematic illustration of transwell-coculture to investigate the effects of cytokines secreted from gastrocnemius and pharyngeal satellite cells. An empty bottom well (Blank) was used as a negative control. B. Representative images of Cell Tracker Green stained gastrocnemius myogenic progenitor cells cocultured with gastrocnemius (GA) or pharyngeal (PH) satellite cells. Scale bars = 330 µm. C. The normalized (ratio to transwell of Blank bottom) cell numbers of gastrocnemius myogenic progenitor cells after Cell Tracker image analysis using cytokine interphase system. n = 3. Statistical significance was determined by 1-way ANOVA. D. Relative mRNA expression level of *Col5a1* in gastrocnemius (GA) and pharyngeal (PH) myogenic progenitor cells (MPCs) and muscles obtained from 3 month old mice. n = 3. Statistical significance was determined by Student’s t-test. E. Representative image of co-localization or adjacent expression of COLV and PAX7 in gastrocnemius muscle section. Scale bars = 35 µm. F. Quantified graph showing the percentage of COLV^+^ niche per PAX7^+^ cells in gastrocnemius (GA), thyropharyngeus (TP) and cricopharyngeus (CP) muscles of *Pax7 Cre*^*ERT*^*-tdTomato* mouse. Statistical significance was determined by 1-way ANOVA. For all graphs, the value represents mean ± SEM. Asterisks indicate statistical significance (*p<0.05, **p<0.01).

### 3.5. Pharyngeal satellite cells are less associated with collagen V, a marker of the quiescent niche

Although we demonstrated that cell-intrinsic factors are at least partly responsible for the increased proliferation and differentiation of pharyngeal SCs, we hypothesized that niche factors including the extracellular matrix (ECM) and other cell types also contribute to the unique phenotypes of pharyngeal SCs. The skeletal muscle extracellular matrix includes several types of collagen and contributes to muscle contraction and maintenance as well as the satellite cell niche (Gillies & Lieber, 2011). A recent study reported that extracellular matrix collagen V (COLV) is secreted by SCs and is important for maintenance of the quiescent state (Baghdadi et al., 2018). We detected decreased levels of the *Col5a1* mRNA, which encodes COLV, in pharyngeal MPCs and muscles relative to gastrocnemius counterparts (Fig 4D). To confirm that reduced *Col5a1* mRNA in pharyngeal muscles is indeed associated with a less quiescent niche, we examined the presence of Collagen V protein relative to PAX7^+^ SCs in pharyngeal and gastrocnemius muscle sections from *Pax7 Cre*^*ERT2*^*-tdTomato* mice. We quantified PAX7^+^ cells within a collagen V^+^ niche, which is defined by co-localization or adjacent location of COLV and PAX7 (Fig 4E). The percentage of PAX7^+^ SCs in the collagen V^+^ niche is significantly higher in GA muscles than in TP and CP muscles (Fig 4F), indicating that pharyngeal SC niches contain less collagen V than limb SC niches. Taken together, these data suggest that the SC niche in pharyngeal muscles is less supportive of the quiescent state than the SC niche in limb muscles.

### 3.5. The pharyngeal muscle niche contains multiple cell types and secreted factors to enhance satellite cell activation

To explore the contribution of extrinsic secreted factors to pharyngeal SC activation, we measured the level of known SC activating factors in pharyngeal muscles. We focused on secreted factors known to modulate SC proliferation including hepatocyte growth factor (HGF), a well-known activator of quiescent SCs upon muscle injury (Allen, Sheehan, Taylor, Kendall, & Rice, 1995), and follistatin (FST), also known as a myostatin inhibitor (Amthor et al., 2004), which induces SC activation and fusion (Gilson et al., 2009; Jones et al., 2015). Interestingly, the mRNA levels of *Hgf* and *Fst* were increased in pharyngeal muscles compared to gastrocnemius muscles (Fig 5A). This result led us to hypothesize that other cell types within pharyngeal muscles contribute to SC activation. Resident macrophages and FAPs have been known to be a primary source of HGF (Sisson et al., 2009) and follistatin (Madaro, Mozzetta, Biferali, & Proietti, 2019), respectively. Thus, we chose these cell types for further analysis. To identify resident macrophages and FAPs in pharyngeal muscles, we stained sections for CD206, a surface marker of resident macrophages (Kosmac et al., 2018) and PDGFRα, a surface marker of FAPs (Joe et al., 2010) (Fig 5B and 5D). The number of CD206^+^ cells per 100 fibers in pharyngeal muscles was significantly higher than in uninjured gastrocnemius (Fig 5C). We also detected a significant increase in PDGFRα ^+^ cells per 100 fibers or per PAX7^+^ cells in pharyngeal muscles relative to uninjured gastrocnemius muscles (Fig 5E and 5F). Although the number of CD206^+^ cells or PDGFRα ^+^ cells in pharyngeal muscles was higher than in uninjured muscles, these numbers were less than the number of CD206^+^ cells or PDGFRα ^+^ cells in 3-day injured limb muscles. Based on these findings, we suggest that the increased numbers of macrophages and FAPs in pharyngeal muscles may activate pharyngeal SCs via secretion of HGF and follistatin.

**Figure 5.**
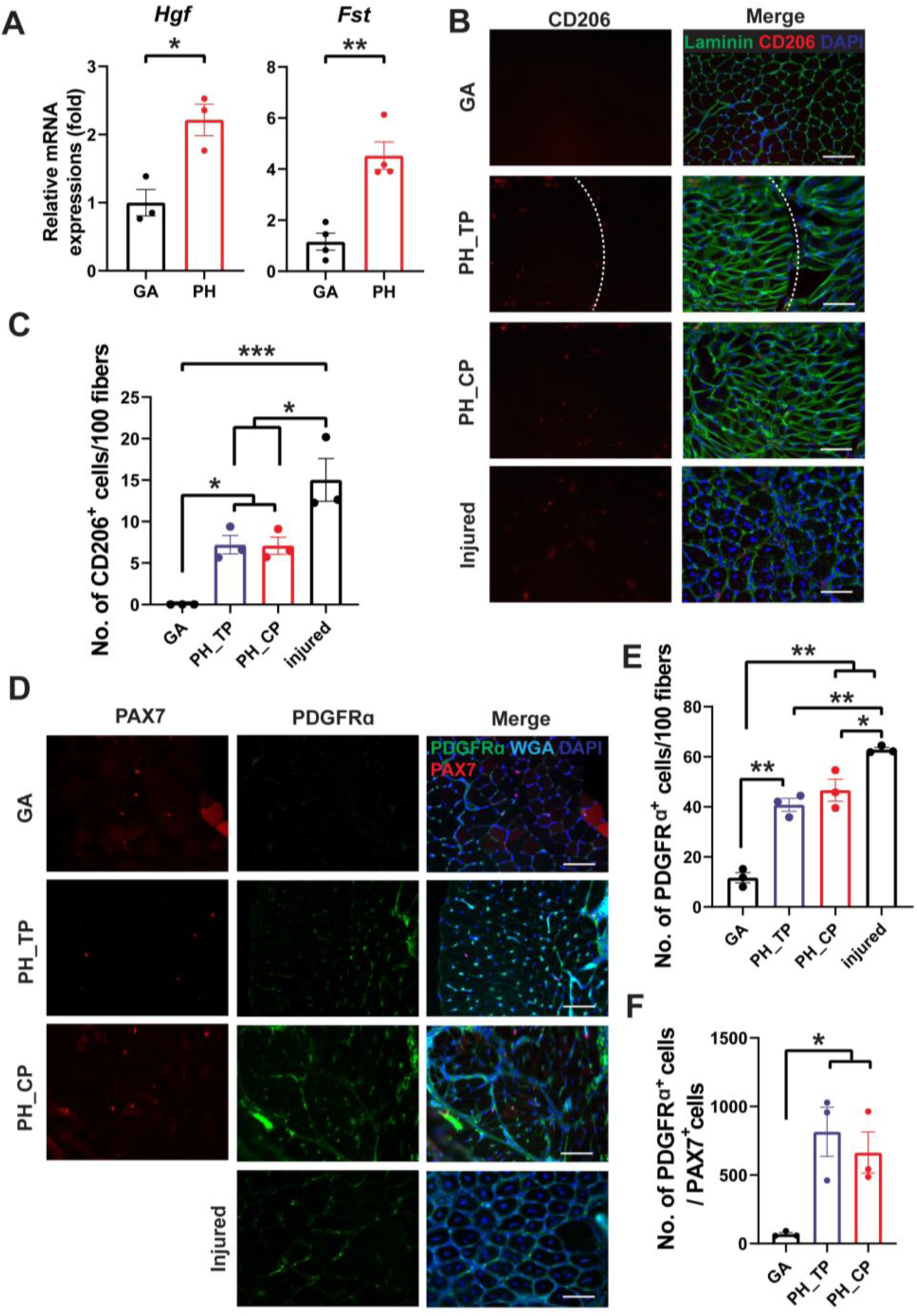
Pharyngeal muscles contain an increased number of resident macrophages and fibroadipogenic progenitor cells (FAPs). A. Relative mRNA expression level of hepatocyte growth factor (*Hgf)* and follistatin (*Fst)* in gastrocnemius (GA) and pharyngeal (PH) muscles obtained from 3 month mice. n = 3. Statistical significance was determined by Student’s t-test. B. Representative images of CD206^+^ cells in gastrocnemius (GA) and pharyngeal (PH) muscles. Merged images show immunostaining with anti-CD206 (Red) and anti-laminin (green) antibodies and DAPI (blue). The asterisk in the right side of dashed white line indicate thyropharyngeus (TP) muscle area. Scale bars = 330 µm. C. Quantified number of CD206^+^ cells per 100 myofibers. BaCl2-injured tibialis anterior (TA) muscles are used as a positive control. n = 3. Statistical significance was determined by ANOVA. D. Representative images of fibroadipogenic progenitors (FAPs) in gastrocnemius (GA) and pharyngeal (PH) muscles. Merged images show immunostaining with anti-PDGFRα (Cyan) and anti-laminin (green) antibodies and DAPI (blue). Scale bars = 330 µm. E. Quantified number of PDGFRα ^+^ cells per 100 myofibers. BaCl2-injured tibialis anterior (TA) muscles are included as a positive control. n = 3. Statistical significance was determined by 1-way ANOVA. F. Quantified number of PDGFRα ^+^ cells per PAX7^+^ cells. n = 3. For all graphs, the value represents mean ± SEM. Asterisks indicate statistical significance (*p<0.05 and **p<0.01).

### 3.6. Pharyngeal satellite cells contribute to muscle homeostasis and niche maintenance of pharyngeal muscle

Given the increased proliferation and differentiation of pharyngeal SCs, we hypothesized that pharyngeal SCs are important for maintaining pharyngeal muscle homeostasis. In a previous study, ablation of SCs in mice from 2 to 6 months of age led to no change of laryngeal pharyngeal muscle fiber cross-sectional area (CSA) but a small significant change of nasal pharyngeal muscle (Randolph et al., 2015). However, in 12 month-old mice, pharyngeal SCs showed significantly higher proliferation (Supplementary Figure 1, (Randolph et al., 2015)) and fusion with consistent SC number compared to pharyngeal SCs of 3 month-old mice (Supplementary Figure 2). Therefore, we investigated the functional importance of pharyngeal SCs around 12 month-old mice by utilizing *Pax7 Cre*^*ERT2*^ *-DTA* mice, which express tamoxifen-inducible, SC-specific diphtheria toxin and thus induce ablation of PAX7^+^ SCs cells *in vivo.* To deplete SC in middle aged mice, we treated with tamoxifen to induce SC-specific DTA expression in 6 month-old mice and harvested pharyngeal muscles after 9 months of SC ablation (Fig 6A). As shown in Fig 6B, we detected a significant decrease in the level of *Pax7* mRNA in the SC-ablated *Pax7 Cre*^*+/-*^*-DTA*^*+/+*^ TM (Tamoxifen) group compared to the control *Pax7 Cre*^*+/-*^*-DTA*^*+/+*^ CO (Corn oil vehicle control) group. There was no significant difference between the *Pax7 Cre*^*+/-*^*-DTA*^*+/+*^ TM (Tamoxifen) group and *Pax7 Cre*^*-/-*^*-DTA*^*+/+*^ TM (Cre control) group. To test whether ablation of PAX7^+^ SCs impacts pharyngeal myofiber size, we stained sectioned TP and CP muscles sections with hematoxylin/eosin and measured the CSA of myofibers. There was no significant change in myofiber CSA in TP muscles of SC-ablated *Pax7 Cre*^*+/-*^*-DTA*^*+/+*^ TM mice relative to controls (Fig 6C). Unexpectedly, the frequency distribution of CSA shifted to the right in CP muscles of *Pax7 Cre*^*+/-*^*-DTA*^*+/+*^ TM mice, indicating a preponderance of larger myofibers after ablation of SCs (Fig 6C). This result implies that SCs in the CP muscle are involved in the maintenance of pharyngeal muscle size.

**Figure 6.**
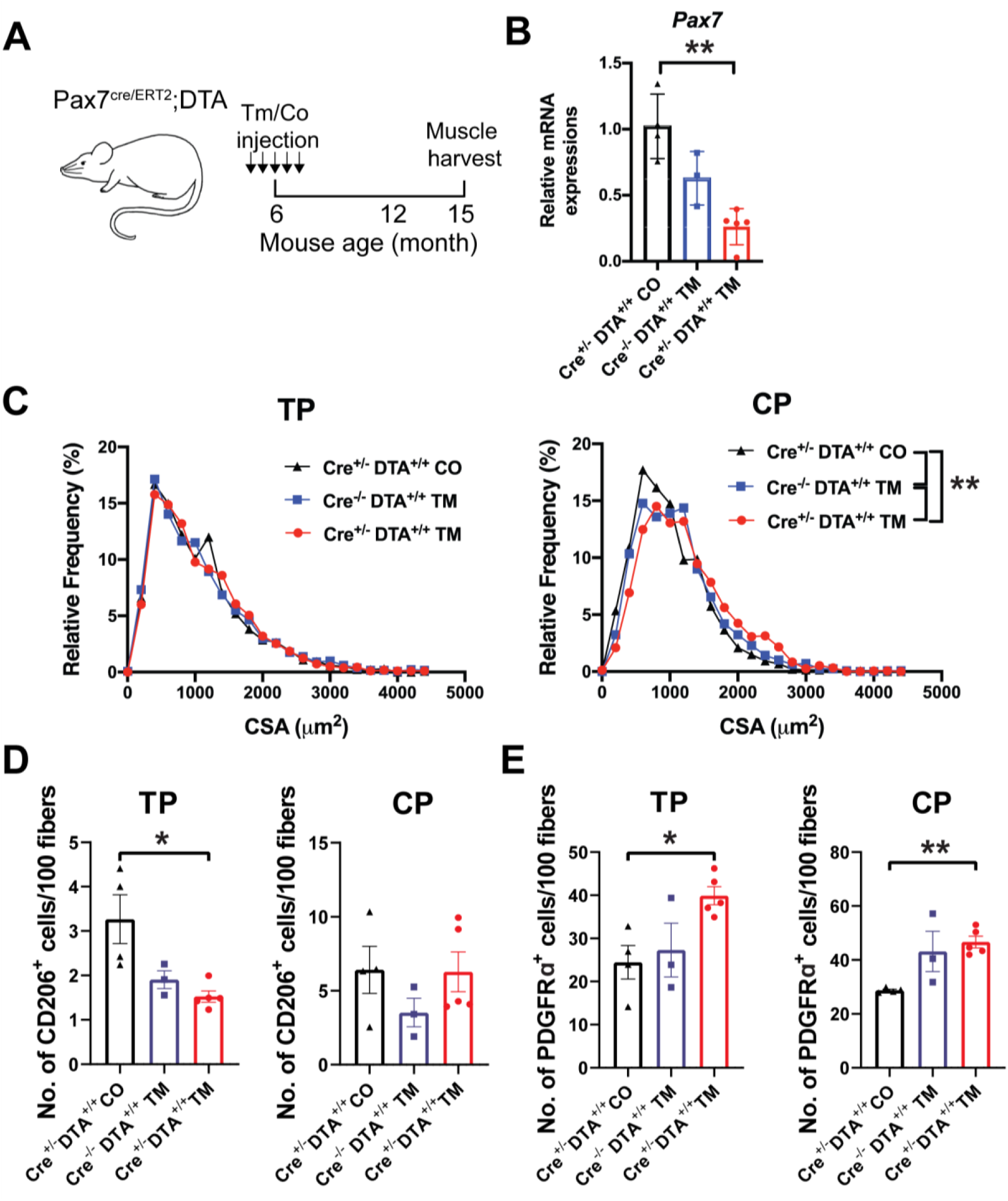
Myofiber cross-sectional area and niche factors are impacted by satellite cell ablation in pharyngeal muscle. A. Scheme of experiments using *Pax7 Cre*^*ERT2*^*-DTA* mice. Abbreviations: Tamoxifen (TM); Control (CO). B. Relative mRNA expression level of *Pax7* in muscle samples (*Pax7 Cre*^*+/-*^;*DTA*^*+/+*^ TM, *Pax7 Cre*^*-/-*^;*DTA*^*+/+*^ TM, *Pax7 Cre*^*+/-*^;*DTA*^*+/+*^ CO) obtained from *Pax7 Cre*^*ERT2*^ *-DTA* mice with or without tamoxifen (TM) treatment. n = 4 or 5. Statistical significance was determined by 1-way ANOVA. C. Frequency distribution plots of myofiber cross-sectional area (CSA) from the thyropharyngeus (TP) and cricopharyngeus (CP) muscle regions are shown. n = 4, with 1400-1800 myofibers combined per condition. Statistical significance was determined by Kruskal-Wallis test. D, E. Quantified number of CD206^+^ cells (D) and PDGFRα ^+^ cells (E) per 100 myofibers in TP and CP of *Pax7 Cre*^*ERT2*^*-DTA* mice. n = 4 or 5. Statistical significance was determined by 1-way ANOVA. For all graphs, the value represents mean ± SEM. Asterisks indicate statistical significance (*p<0.05 and **p<0.01).

Considering that SCs are known to signal to other cell types in the skeletal muscle niche (Fry, Kirby, Kosmac, McCarthy, & Peterson, 2017), we hypothesized that loss of secreted factors from pharyngeal SCs affect the neighboring cells close to the SC niche. We sectioned pharyngeal muscles from SC-ablated *Pax7 Cre*^*ERT2*^ *-DTA* mice and quantified CD206^+^ cells (resident macrophages) and PDGFRα ^+^ cells (FAPs) using immunofluorescent staining and microscopy (Fig 6D-E). We observed significantly decreased numbers of CD206^+^ cells in the TP muscle of *Pax7 Cre*^*ERT2*^ *-DTA* mice (Fig 6D). Conversely, we detected a significant increase in the number of PDGFRα^+^ cells in the TP and CP muscles of *Pax7 Cre*^*ERT2*^ *-DTA* mice (Fig 6E). Although the difference was not statistically significant, *Cre*^*-/-*^*-DTA*^*+/+*^ TM (Cre control) mice showed a trend of reduced *Pax7* expression (Fig 6B) as well as similar trends of neighboring cell pool change with *Pax7 Cre*^*ERT2*^ *-DTA* mice (Fig 6D-E). Taken together, these data suggest that pharyngeal SCs are required not only to maintain the size of myofibers in cricopharyngeus muscles but also to control the number of neighboring macrophages and FAPs that contribute to the microenvironment of the pharyngeal SC niche.

## 4. Discussion

Although craniofacial muscles including pharyngeal muscles differ from body muscles in embryonic origin and core genetic programs (S Tajbakhsh, 2009), the majority of satellite cell (SC) studies have focused on those in the limb muscles. Studies of SCs in craniofacial muscle are difficult due to its small size, the difficulty dissection, and the lack of functional assays. To overcome these difficulties, we employed a series of genetic mouse models to either label or ablate SCs and reveal the distinct characteristics of pharyngeal muscle SCs. To explain the highly proliferative and differentiative properties of pharyngeal SCs, we investigated both intrinsic factors, such as mitochondria and autocrine factors, and extrinsic niche including ECM and secreted factors from other cell types within pharyngeal muscles. Our study provides new evidence to explain how both intrinsic mechanisms and extrinsic factors govern the unique state of craniofacial SCs.

### G-alerted features of pharyngeal SCs: cell size and mitochondria contents

Dividing SCs typically follow one of two fates, either a return to the quiescent G_0_ state to renew the SC pool (Li & Clevers, 2010) or entry into an actively cycling G_1_ state to proceed along the myogenic lineage (Yin, Price, & Rudnicki, 2013). A third state, G_Alert_, has also been described, which is intermediate to quiescence and activation (Rodgers et al., 2014). Our data indicate that SCs in uninjured pharyngeal muscles exhibit multiple aspects of the G_Alert_ state including increased cell size, enhanced mitochondrial mass, and increased propensity to proliferate and differentiate (Rodgers et al., 2014). Given that SCs in the G_Alert_ state are ‘primed’ to rapidly respond to injury stimuli, the G_Alert_-like state of pharyngeal SCs suggests a poor ability to maintain quiescence and, once activated, their cell fate is more forced toward commitment when compared to quiescent limb SCs. Indeed, we detected enhanced proliferation and differentiation of cultured SCs isolated from pharyngeal muscle relative to those isolated from the gastrocnemius. This unique “alert”-like state is thought to be tightly regulated by distinct intrinsic or extrinsic factors of pharyngeal muscles itself: cellular metabolism, cell autonomous signaling, cell-cell signaling, the extracellular environment, inflammatory mediators, and so on (Aurora & Olson, 2014; Quarta et al., 2016).

One of the core intrinsic factors governing SC state is mitochondria-mediated metabolic regulation, which is considered critical for cell fate decisions, activation, and myoblast proliferation (Duguez, Sabido, & Freyssenet, 2004; Zhang, Menzies, & Auwerx, 2018). We found that pharyngeal SCs showed relatively higher mitochondrial content relative to quiescent limb SCs, which contain relatively few mitochondria. This result is consistent with a previously reported increase in mitochondrial content upon SC activation in limb muscle, despite a shift to glycolytic metabolism (Montarras, L’honoré, & Buckingham, 2013; Ryall et al., 2015). Interestingly, impaired mitochondrial function has been associated with the pathology of oculopharyngeal muscular dystrophy, which preferentially affects craniofacial muscles (Chartier et al., 2015; Vest et al., 2017). Further investigation is necessary to investigate the importance of mitochondrial metabolism in pharyngeal SC biology and in pharyngeal muscle pathologies.

### Pharyngeal SC-derived niches: auto/paracrine factors and collagen V

The skeletal muscle niche is critically important in regulating SC state. We tested two important components of the SC-derived niche including autocrine signaling via secreted factors and ECM components. In co-culture experiments, we determined that pharyngeal SCs enhance proliferation of limb MPCs via secreted factors, though further studies are needed to determine the identity of the soluble signals secreted by pharyngeal SCs. Our study also shows that the pharyngeal muscle niche contains less collagen V protein, which is produced by SCs and is considered to be a dominant regulator of the quiescent niche in skeletal muscles (Baghdadi et al., 2018). This result is consistent with the differentially expressed ECM genes pharyngeal SCs reported in a previous microarray study (Randolph et al., 2015). It is possible that other proteoglycans or ECM proteins contribute to regulation of quiescence in a pharyngeal SC-specific manner. Taken together, these data add weight to the hypothesis that pharyngeal SCs are intrinsically less quiescent.

### Neighboring cells to activate pharyngeal SCs: resident macrophages and FAPs

Neighboring cells contribute to the microenvironment of SCs via soluble factors or direct cell-to-cell contact. HGF is one such auto/paracrine factor involved in SC activation in response to muscle injury, overuse, or mechanical stretches (Miller, Thaloor, Matteson, & Pavlath, 2000; Sheehan, Tatsumi, Temm-Grove, & Allen, 2000; Tatsumi, 2010; Tatsumi, Anderson, Nevoret, Halevy, & Allen, 1998). HGF is secreted into the extracellular matrix of uninjured muscles as pro-HGF and, upon injury, proteolysis by urokinase-type plasminogen activator (uPA) (Sisson et al., 2009) or HGF activator (Rodgers, Schroeder, Ma, & Rando, 2017) generates active HGF that in turn activates SCs (Bernet-Camard, Coconnier, Hudault, & Servin, 1996; Sisson et al., 2009; Stoker, Gherardi, Perryman, & Gray, 1987). Although we did not determine which form of HGF is found in pharyngeal muscle, previous microarray data comparing pharyngeal SCs with limb SCs (Randolph *et al.*, 2015) revealed increased levels of the *Plat* gene encoding tissue-type plasminogen activator (tPA), which is similar (identity 32.8% and similarity 43%) to uPA and cleaves pro-HGF (Mars, Zarnegar, & Michalopoulos, 1993). Thus, pharyngeal SCs may contribute to processing pro-HGF to the active form without injury. Interestingly, extraocular muscle SCs, which also proliferate and differentiate without injury, contain high levels of *Plat* mRNA relative to limb SCs (Pacheco-Pinedo et al., 2009). Thus, HGF is likely an important signal modulating pharyngeal and extraocular SC activity, but additional studies are needed to better define the mechanism.

Although HGF signaling can occur in an autocrine manner in SCs, the majority of HGF is secreted by macrophages (Sisson et al., 2009) that infiltrate the skeletal muscle niche after injury (Pillon, Bilan, Fink, & Klip, 2013). Previous microarray analysis revealed that pharyngeal SCs contain elevated levels of mRNAs encoding cytokines known to attract macrophages (*Ccl2, Ccl12, Ccl7)* or induce macrophage polarization (*Lif* and *Il-6*) (Duluc et al., 2007; Xuan, Qu, Zheng, Xiong, & Fan, 2015). As expected, we detected relatively high numbers resident macrophage without injury in pharyngeal muscles compared to uninjured limb muscles. To further address the role of SCs in macrophage recruitment to pharyngeal muscles, we utilized a genetic mouse model (*Pax7 Cre*^*ERT2*^ *-DTA* mice) to ablate satellite cells. We found that SC ablation led to a decrease in the number of M2 macrophages in the thyropharyngeal muscle. This result confirms that pharyngeal SCs are important for recruiting resident M2 cells under basal conditions. However, the roles of M2 for pharyngeal SC activation as well as function of pharyngeal muscles remain to be determined.

Another niche signal important for modulating SC activity is follistatin, which is a TGF-β antagonist (Amthor et al., 2004) suggested to prime myoblasts for myogenic differentiation and promote myofiber hyperplasia (Jones et al., 2015; Medeiros, Phelps, Fuentes, & Bradley, 2009). We provide evidence that pharyngeal muscles have the increased levels of follistatin mRNA and its cellular source, FAPs. (Reggio et al., 2020). These results are consistent with previous reports in extraocular muscles (Formicola et al., 2014) and in contralateral limb muscles, which contains G_Alert_ SCs (Rodgers et al., 2014). Interestingly, we found that SC may regulate the pool of FAPs, as we detected elevated numbers of FAPs in SC-depleted muscles, which was consistent with previous report using limb muscles (Formicola et al., 2018). However, fibrosis and fatty infiltration were not detected in SC-depleted pharyngeal muscles unlike chronic injured muscles (X. Liu et al., 2016). In contrast, mice in which the FAPs population has been ablated showed reduced numbers of SC in limb muscles indicating that FAPs are important for maintaining SC pools (Ancel, Mashinchian, & Feige, 2019; Wosczyna et al., 2019). Taken together, our results and previously published studies indicate that correlations exists between SC and FAPs pools in skeletal muscle.

### Role of pharyngeal SC for muscle maintenance

To understand how SCs participate in pharyngeal muscle maintenance, we measured the myofiber CSA in cricopharyngeal and thyropharyngeal muscles in SC-ablated mice. Unlike other SC-depletion studies (Keefe et al., 2015; Pawlikowski, Pulliam, Dalla Betta, Kardon, & Olwin, 2015; Randolph & Pavlath, 2015), the conditional depletion of pharyngeal SCs from adult mice led to an increase in the CSA of cricopharyngeal myofibers. These results indicate that SCs are required for maintenance of cricopharyngeus muscle size, though the mechanism allowing for this adjustment remains completely unknown. However, it is inconsistent with our expectation and the previous report showing that satellite cell ablation led to reduced CSA in 12-month SC-depleted EOM (Keefe et al., 2015) and in 4-month SC-depleted nasopharyngeal muscles (no CSA change in laryngopharynx, which contains our target muscles, CP and TP) (Randolph & Pavlath, 2015). One possible explanation for our findings is that the ablation of SCs is incomplete and the remaining diminished population of SCs in pharyngeal muscles produces a hypertrophic response related to the increase in the number of FAPs. Alternatively, hypertrophy of pharyngeal muscle may be a result of small fiber fusion in SC-depleted conditions or SC-independent hypertrophic growth (McCarthy et al., 2011).

In conclusion, this study is the first comprehensive analysis of both intrinsic and extrinsic factors associated with the highly proliferative and differentiative features of pharyngeal SC. While it is not clear whether the unique embryonic origins of pharyngeal SCs leads intrinsic differences in proliferative and myogenic properties, this study demonstrates that both pharyngeal SC-secreted factors and pharyngeal muscle niches are capable of activating pharyngeal SCs without injury. Although the role of highly active SCs in pharyngeal muscle function is still ambiguous, we propose that unique properties of pharyngeal SCs are a promising area of study to better understand pharyngeal muscle specific pathologies, such as oculopharyngeal muscular dystrophy.

---

**Supplementary Figure 1.**
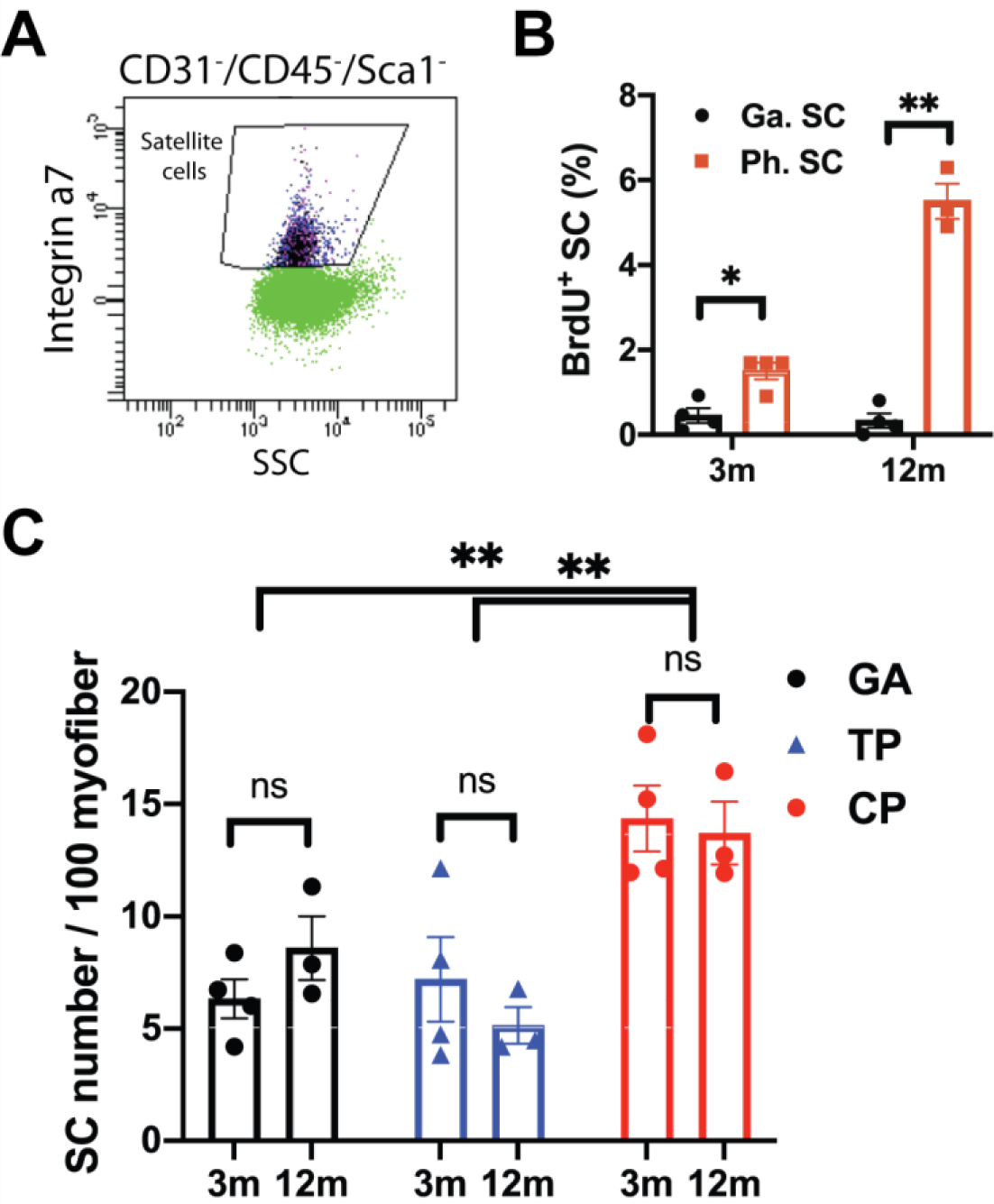
Active *in vivo* proliferation of pharyngeal satellite cells of 12 months old mice compared with ones of 3 months old mice. A. Flow cytometry gating strategy for satellite cells defined by CD31^-^/CD45^-^/Sca1^-^/Intergrin α7^+^. B. Percentage of BrdU^+^ SCs in gastrocnemius and pharyngeal muscles of 3 and 12 month old mice. n = 4 for each age group. Statistical significance was determined by 2-way ANOVA. C. Number of SC in gastrocnemius (GA), thyropharyngeus (TP) and cricopharyngeus (CP) muscles of 3 and 12 month old mice. n = 3 or 4 for each age groups. The value represents mean ± SEM. Statistical significances was determined by 2-way ANOVA. Asterisks indicate statistical significance (*p<0.05 and **p<0.01).

**Supplementary Figure 2.**
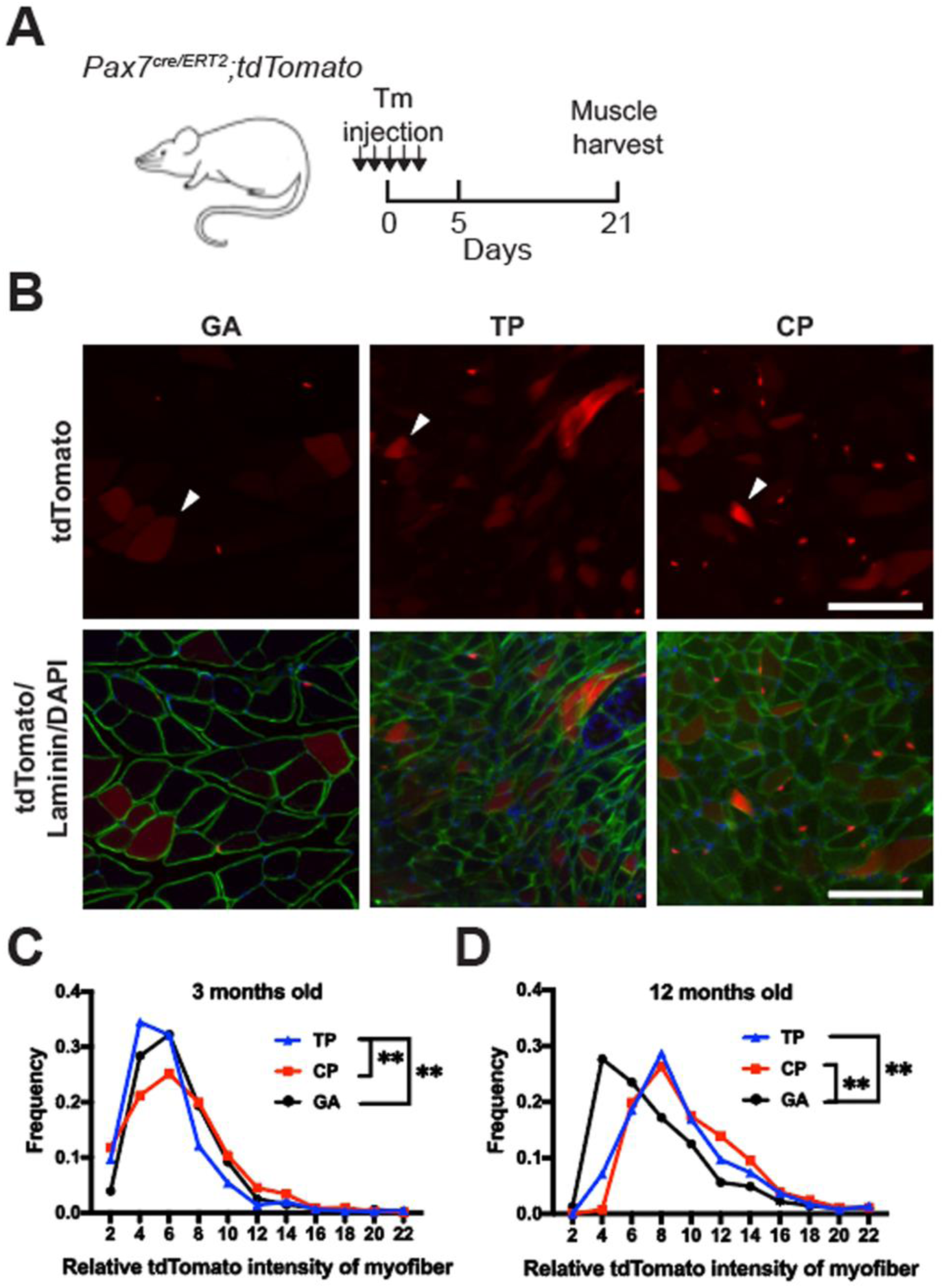
Increased fusion of pharyngeal satellite cells in 12 month old mice compared with 3 month old mice. A. Scheme of experiments using *Pax7 Cre*^*ERT2*^*-tdTomato* mice. Abbreviations: Tamoxifen (TM); Control (CO). B. Representative cross-section expressing PAX7^+^ (red) satellite cells in gastrocnemius (GA), thyropharyngeus (TP) and cricopharyngeus (CP) muscles from 12 month old *Pax7 Cre*^*ERT2*^*-tdTomato* mice. White arrow heads indicate examples of PAX7^+^ SC fused myofibers, which show diffused tdTomato expression inside the myofiber. Basal lamina was immune-stained with anti-Laminin antibody (green), and DAPI (blue) was used to stain nuclei. C, D. Frequency distribution of tdTomato intensity in myofibers from gastrocnemius (GA), thyropharyngeus (TP) and cricopharyngeus (CP) muscles of 3 month (C) and 12 month (D) old *Pax7 Cre*^*ERT2*^*-tdTomato* mice. Statistical significance was determined by Kruskal-Wallis test. Asterisks indicate statistical significance (**p<0.01).

**Supplementary Table 1.**
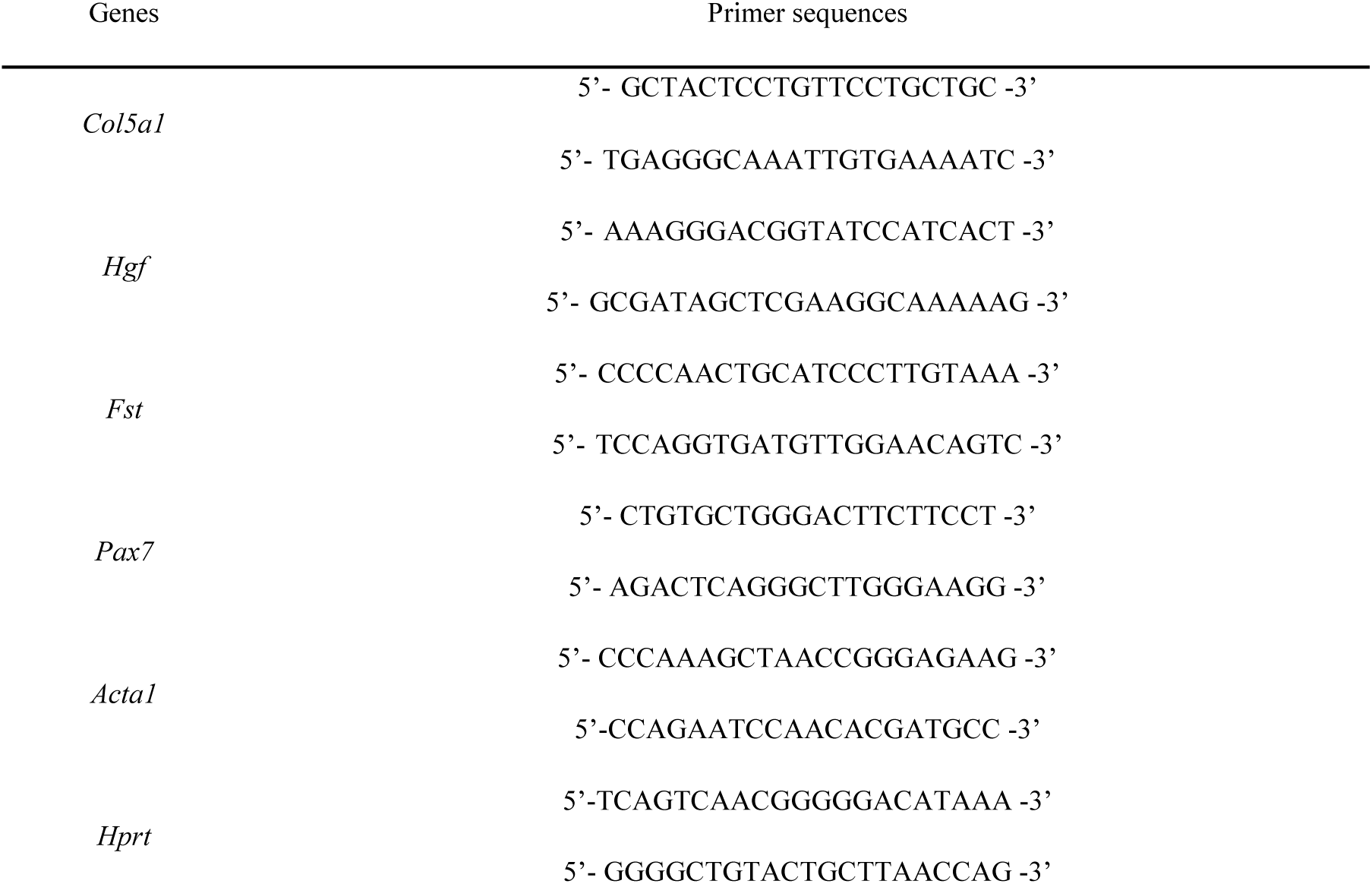
Primers used for gene expression analysis.

**Supplementary Table 2.** Antibodies used for immunofluorescence staining.

| Detection of | Name | Host species | Dilution or Concentration | Manufacturer (Cat #) |
| --- | --- | --- | --- | --- |
| Resident M2 macrophage | Mannose Receptor (CD206) | Rabbit IgG | 1:200 | Abcam (ab125028) |
| Fibroadipogenic progenitors (FAPs) | PDGFRα | Rabbit IgG | 1:200 | Cell Signaling Tech. (3174S) |
| Collagen V A1 | COLVA1 | Rabbit IgG | 1:200 | Sigma (SAB1306996) |
| Basement membranes | Laminin | Rabbit IgG | 1:400 | Sigma (L9393) |
| Basement membranes | Wheat Germ Agglutinin (WGA) | Alexa Fluor™ 647 Conjugate | 1:400 | Invitrogen (W32466) |

## Acknowledgements

This research was supported in part by grants from National Institutes of Health (NIH) NIAMS (R01 AR071397) and Basic Science Research Program through the National Research Foundation of Korea (NRF) funded by the Ministry of Education (NRF-2018R1A6A3A03011703).

## Author contributions

EK designed the study, performed experiments, image analysis, analyzed data, prepared figures and wrote the manuscript. YZ performed experiments and image analysis. FW performed experiments and analyzed data. JA prepared samples. KEV performed experiments, analyzed data and wrote the manuscript. HJC designed the study, performed experiments, image analysis, analyzed data, prepared figures and wrote the manuscript.

## Conflict of interest

The authors declare that they have no conflict of interest.

## Data availability

All data to support the conclusions of this manuscript are included in the main text and supplementary materials.

## Notes

### Competing Interest Statement

The authors have declared no competing interest.

